# Acidification by nitrogen metabolism triggers extracellular biopolymer production in an oleaginous yeast

**DOI:** 10.1101/2025.05.04.652101

**Authors:** Henrique Sepúlveda Del Rio Hamacek, Oksana Tingajeva, Katharina Ostertag, Alīna Reķēna, Aleksandr Illarionov, Piia Jõul, Paola Monteiro de Oliveira, Giselle de La Caridad Martín-Hernández, Bettina Müller, Nemailla Bonturi, Volkmar Passoth, Petri-Jaan Lahtvee, Rahul Kumar

## Abstract

The oleaginous yeast *Rhodotorula toruloides* is a natural producer of lipids and carotenoids. However, its potential as a producer of extracellular biopolymers remains unexplored. Hence, we aimed to evaluate the *R. toruloides* CCT0783 for extracellular biopolymer production. We found that the carbon-to-nitrogen ratio influenced exopolysaccharide (EPS) production, reaching 5.84 ± 0.45 g/L under glucose-rich conditions. We characterized the crude biopolymer using FTIR and GC-MS, identifying polysaccharide peaks, wherein EPS consisted of (%) glucose (88.83 ± 4.87), galactose (5.50 ± 1.50), mannose (4.80 ± 1.57), and xylose and arabinose (0.87 ± 0.04) monomers. The dried EPS also contained a fractional presence of protein (1.0%). Interestingly, inorganic, but not organic, nitrogen metabolism was associated with the acidification of the culture environment and simultaneous EPS production. A comparison between non-buffered and buffered media indicated that EPS was produced when the final pH was around or below 2, which was observed in the former but not in the latter. Finally, a comparative bioinformatics analysis allowed us to map a putative EPS biosynthesis and transport pathways, as well as regulators of intracellular pH maintenance. In conclusion, our study demonstrates *R. toruloides’* potential as an extracellular microbial biopolymer producer, enabling its development as a cell factory.

**Importance:** Microbial biopolymers are extensively investigated for their impact on the environment and health. However, developing a biotechnology process for producing such biopolymers remains challenging despite their potential for valuable applications. Considering this opportunity, we investigated the oleaginous yeast *Rhodotorula toruloides* as a producer of extracellular biopolymers. Our study describes the conditions of exopolysaccharide (EPS) biosynthesis in *R. toruloides*. This extracellular biopolymer could have a wide range of applications, from gelling agents in pharmaceuticals to emulsifiers in the food industry. Furthermore, our comparative bioinformatics analysis results could be useful for metabolic engineering and developing *Rhodotorula* cell factories.

## 1. Introduction

*Rhodotorula toruloides* is a basidiomycete oleaginous yeast known for its natural ability to produce carotenoids and neutral lipids^1^. Additionally, it can be engineered to synthesize value-added chemicals such as fatty-acid esters, fatty alcohols, and terpenes^1,2^. In *R. toruloides*, significant advancements have been made in improving lipid and carotenoid production by optimization of cultivation conditions^3,4^, deploying metabolic engineering tools^5^, or performing laboratory adaptive evolution^6^. For instance, nutrient limitation conditions (nitrogen, phosphorus, and sulfur) increase lipid and carotenoid biosynthesis^4,7^. Salt stress reportedly enhances carotenoid production, wherein slight changes in culture density and a negative impact on glucose metabolism were observed compared to conditions without high salt concentrations^8^.

Due to its ability to grow on a wide range of carbon and nitrogen sources and produce high-value chemicals, it has emerged as an attractive candidate for designing microbial cell factories^9^. Moreover, *R. toruloides’* ability to consume diverse substrates, such as crude glycerol, lignocellulosic hydrolysates, and wastewater, and to tolerate inhibitory compounds in the environment makes it a robust model organism for the development of sustainable processes^10–13^.

In biotechnology, many microbial cell factories are based on yeast fermentation of sugars, producing extracellular biochemicals partially due to the Crabtree effect, where excess carbon overflows in metabolism are directed to fermentation products, allowing energy and redox cofactors maintenance during cellular growth^14,15^. *R. toruloides* is a Crabtree-negative yeast; therefore, aside from carbon dioxide, it does not typically produce a major byproduct metabolite in conditions with high concentrations of sugars^16,17^. However, wild-type *R. toruloides* (strains NRRL Y-1588, NRRL Y-1091, NRRL Y-6672, and NRRL Y-17902) were reported to produce slime on leaf surfaces, where a typical biofilm formation may occur in the native environment^18–20^. Interestingly, in our previous study^8^ on *R. toruloides* CCT0783, we observed cellular aggregate formation in laboratory conditions, which led us to hypothesize that these aggregates may form due to cell-cell interactions, potentially caused by the production of extracellular polymeric substances in this yeast. However, such secreted products have not been previously quantified in *R. toruloides* CCT0783, motivating our present study.

In principle, yeasts, bacteria, algae, and other microorganisms can produce exopolysaccharide (EPS) as a secondary metabolite^20^. The EPS provides those microbes with a stable microenvironment where they can be protected against exogenous stressors such as high temperature variations, low or high pH levels, UV radiation, desiccation, and oxidative compounds^21^. EPS is also known to help yeasts and bacteria adhere to natural and synthetic surfaces through biofilm formation^22^. Biopolymers sourced from microorganisms, including xanthan, dextran, scleroglucan, polyhydroxyalkanoates (PHA), and poly(ester amide)s (PEAs), are highly valuable in industrial applications^23,24^. Microbial polysaccharides exhibit particular chemical and physical properties^21^, enabling their use as emulsifiers, stabilizers, binders, gelling agents, coagulants, lubricants, and thickening agents in the food industry^25^. Pharmaceutical, environmental, and sustainable production of materials are other areas of EPS applications^21^. Therefore, an investigation of EPS production conditions and identification of potential biosynthesis pathways in *R. toruloides* are relevant for its development as a microbial cell factory, and hence, a focus of our study, as EPS might play a crucial role in being used as sustainable polymers with tailorable properties to help meet future demands across various applications.

Though extensive research on lipid and carotenoid production in *R. toruloides* is available, the production of extracellular biopolymers remains relatively unexplored. Because of this, our current study is pertinent for filling the knowledge gap, where we investigated extracellular biopolymer production by this yeast. We used qualitative and quantitative approaches to observe physiological parameters that influence extracellular biopolymer production and characterize the EPS using FTIR and GC-MS techniques. Our results show that most of the extracellular biopolymers produced by *R. toruloides* are polysaccharides, while a minor fraction is protein. For a holistic understanding of the biopolymer production process, we performed a comparative bioinformatics analysis of publicly available datasets for *R. toruloides*, mapping the putative metabolic pathways for EPS biosynthesis and transport. Finally, our results suggest a possible association between the acidification of the media, the utilization of inorganic nitrogen, and biopolymer production in this yeast.

## 2. Materials and Methods

### 2.1. Strain and culture conditions

The yeast *Rhodotorula toruloides* CCT0783 (synonym NBRC10076)^8,26^ is used in this study. The cultures were maintained in glycerol stocks, stored at −80 °C, and aliquoted vials were thawed on ice for inoculation. Pre-culture cultivations were performed in a 50-mL Falcon tube containing 5 mL YPD medium and incubated overnight (200 rpm, 30 °C) in a rotary shaker incubator (Excella E25, New Brunswick Scientific, USA). The obtained cells were then used as the inoculum in cultures with chemically defined media. A consistent inoculum size was achieved by measuring pre-culture optical density at 600 nm (OD_600_) and ensuring the initial OD_600_ was 0.1 after inoculation in experiments. Experiments were performed using 250 mL sterile shaker flasks containing 25 mL of chemically defined media. The flasks were kept in a rotary incubator (200 rpm, 30 °C) for 120 h. Samples for measuring dry cell weight, EPS, and pH values were collected as needed. All preparations and aliquoting were conducted under sterile conditions.

### 2.2. Culture media: composition and preparation

YPD medium was used to revive cells from glycerol stocks and to prepare pre-cultures. One liter of YPD medium contained 20 g of peptone from meat (bacteriological grade, >95%, Thermo Fisher Scientific, USA), 10 g of yeast extract (Thermo Fisher Scientific, USA), and 22 g of D(+)-glucose monohydrate (>99.5%, Carl Roth GmbH, Germany). The YPD medium had its pH adjusted to 3 with a solution of HCl (1 mol/L) when needed.

The chemically defined culture media were prepared with Milli-Q water according to formulations described previously^8^, with the following composition, per liter: 3 g KH_2_PO_4_ (100%, Thermo Fisher Scientific, USA); 2 or 5 g (NH_4_)_2_SO_4_ (99.8%, Thermo Fisher Scientific, USA); 0.5 g MgSO_4_•7H_2_O (≥99%, Thermo Fisher Scientific, USA); 20 to 120 g D(+)-glucose monohydrate or ethanol (96% v/v). The pH was adjusted to 3 with an HCl solution (1 mol/L) when needed. K_2_HPO_4_ (98%, 5.25 g/L) was added to give a pH of approximately 6.9 for the buffered cultivation media. Solid KCl was added to the cultivation media in specific experiments to obtain a concentration of K^+^ ions equal to 0.1 mol/L. Vitamins and trace elements solutions were added in the proportion of 1 mL per liter of medium. They were prepared and sterilized by filtration separately. The composition of the vitamin solution per liter of Milli-Q water (pH = 6.5) was biotin, 0.05 g; p-amino benzoic acid, 0.2 g; nicotinic acid, 1 g; Ca-pantothenate, 1 g; pyridoxine-HCl, 1 g; thiamine-HCl, 1 g; and myoinositol (AppliChem GmbH, Germany), 25 g. The composition of the trace elements solution per liter of Milli-Q water was EDTA, 15 g; ZnSO_4_•7H_2_O, 4.5 g; MnCl_2_•2H_2_O, 0.84 g (or MnCl_2_•4H_2_O, 1.03 g); CoCl_2_•6H_2_O, 0.3 g; CuSO_4_•5H_2_O, 0.3 g; Na_2_MoO_4_•2H_2_O, 0.4; H_3_BO_3_, 1 g; KI, 0.10 g^15^.

Thermal sterilization of the media components was conducted in an autoclave (Systec V-95, Systec, USA) at 121 °C for 15 minutes. After the solutions were cooled to room temperature, they were mixed, and the vitamins and trace elements were added. All steps were performed in a sterile environment. Experimental conditions were varied by changing the C:N ratio (Equation 1), in which N_c_ and N_N_ represent the number of moles of carbon and nitrogen atoms per mole of the corresponding carbon and nitrogen sources; [C-source] and [N-source] represent the concentrations of the carbon and nitrogen sources (mol/L), respectively; and MM_C_ and MM_N_ represent the molar masses (g/mol) of the carbon and nitrogen sources, respectively.

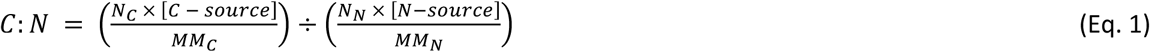

### 2.3. High-Performance Liquid Chromatography (HPLC)

To obtain the initial concentration of glucose and ethanol in the media, 1 mL of medium was collected in 2-mL Eppendorf^®^ tubes, centrifuged (21950 × *g*, 4 °C, 5 min), and the supernatant was subsequently transferred to HPLC vials. The samples were filtered and diluted as necessary. Glucose and ethanol in the solution were quantified using an HPLC (LC-2030C Plus, Shimadzu, Kyoto, Japan) equipped with a refractive index detector (RID-20A, Shimadzu, Kyoto, Japan). The samples were stored at 4 °C on the sample rack, and 20 μL aliquots were automatically injected into an Aminex HPX-87H ion exclusion column (Bio-Rad, US). Elution with 5 mmol/L H_2_SO_4_ was performed in isocratic mode at 0.6 mL/min with column and detector temperatures set to 45 °C.

### 2.4. pH measurements

To measure the initial pH value, 1 mL of the culture medium was transferred to 2 mL Eppendorf^®^ tubes, and the pH was determined using a pH meter (HACH HQ30d, METTLER TOLEDO, USA). For extra pH measurements at specified cultivation times, the same volume was centrifuged (21950 × *g*, 20 °C, 5 min) to pellet the cells. Then, the supernatant was collected for pH measurement.

### 2.5. Determination of dry cell weight

The biomass dry cell weight (DCW) was obtained by collecting 1 mL of cultivation medium and then centrifuging it (21950 × *g*, 5 min). The supernatant was removed, and 1 mL of PBS was added to resuspend the cells. The samples were vortexed, pipetted into pre-weighted 0.45 μm filter papers, and filtered using a vacuum filtration system. The filter papers containing the biomass were dried in a microwave oven at 900 W for 2-3 minutes in 30-second intervals. The filter papers were stored in a desiccator until weighed the following day. The difference in the measured weight represented the DCW of the samples.

### 2.6. Extracellular Biopolymers: Polysaccharides and Proteins

#### 2.6.1. Extraction and quantification

Following the procedures described in the literature^27,28^, the exopolysaccharides (EPS) were separated from the cell culture with minor modifications. Briefly, the culture medium was transferred to 50-mL Falcon tubes and centrifuged (6000 × *g*, 4 °C, 30 min). The supernatant was carefully separated from the cell pellet. To precipitate the EPS, cold ethanol (96% v/v, 4 °C) was added to the supernatant at a ratio of 2 parts of ethanol to 1 part of supernatant in volume. The mixture was stored at 4 °C for 24 h to promote EPS precipitation. After this period, a distinct layer of EPS formed at the middle or top of the liquid. This layer was carefully collected for further processing and analysis. Finally, the obtained EPS was placed in an incubator at 50 °C until no change in mass was observed, indicating that all the excess liquid had evaporated. The obtained product was qualitatively confirmed to be a polysaccharide through the Alcian blue staining method with minor modifications^29^. In brief, a stock solution was prepared by dissolving 1% w/v Alcian Blue 8GX in 3% v/v acetic acid solution. The cell suspension was stained for 30 minutes in a diluted staining solution with a final concentration of 0.08% w/v Alcian Blue 8GX. The stained cell suspension was observed under an Olympus Microscope CX21 (Olympus, Japan).

Protein quantification of dried EPS samples was performed by Bradford’s assay^30^ with Pierce^TM^ Bradford Protein Assay Reagent (Thermo Fisher Scientific, USA), following the supplier’s protocol for standard test tube procedures. Briefly, a known mass of dried EPS was dissolved in saline solution (0.9% w/v). Then, 30 μL of the samples or the standards were added to a cuvette containing 1.5 mL of the Bradford reagent. The cuvettes were incubated in the dark and at room temperature for 10 minutes. Then, the absorbance was measured at 595 nm. A bovine serum albumin (BSA) stock solution (2 mg/mL) was used as the standard to prepare the calibration curve following the supplier’s instructions.

#### 2.6.2. Chemical characterizations

##### 2.6.2.1. Fourier Transform Infrared (FTIR) Spectroscopy analysis

For FTIR analysis, dried EPS samples were carefully placed on the ATR crystal and analyzed on an FTIR spectrometer (IRTracer-100, Shimadzu, Kyoto, Japan) in attenuated total reflection (ATR) mode. The spectra were recorded in the range of 4000 to 400 cm⁻¹, with a spectral resolution of 6 cm⁻¹ and an aperture of 5 mm. The peaks identified by the software (LabSolutions, Shimadzu, Japan) were compared to a reference library to determine the functional groups present in the polysaccharide.

##### 2.6.2.2. EPS composition

The monosaccharide composition of the EPS was determined using methods adapted from Hamidi et al.^31^. In brief, 10 mg of EPS were dissolved and hydrolyzed in 1 mL of trifluoroacetic acid (TFA, 2 mol/L) for 90 minutes in a dry bath at 120 °C. The TFA was then evaporated under a nitrogen stream. One mL of sodium borohydride solution (NaBH_4_, 20 g/L) was added, and the reaction was maintained at room temperature for 3 h. The sample was dried again with nitrogen gas. For derivatization, the sample was treated with pyridine (300 μL) and acetic anhydride (400 μL) as described by Wang et al.^32^. The resulting mixture was kept at room temperature and heated to 90 °C for 30 minutes under slow agitation. Then, the sample was centrifuged, and the supernatant was transferred to a new glass vial to separate the solids.

The clarified samples were transferred to a 7890A gas chromatograph coupled to a 5975C mass spectrometer (Agilent Technologies, USA) with an electron ionization source and a quadrupole mass analyzer. One μL of the sample was injected in split mode (1:10) at 275 °C. The flow rate of the carrier gas (helium) was 1.3 mL/min. The initial oven temperature was set to 150 °C (1 min), increasing by 10 °C/min until it reached 220 °C (5 min), and increasing by 10 °C/min until it reached 250 °C (2 min). The total running time was 28 minutes. The analyte ionization was performed in electron ionization mode with an electron energy of 70 eV. The interface, ion source, and mass analyzer temperatures were set at 280, 230, and 150 °C, respectively. Scan mode in the target ion range, from 50 to 800 m/z, was employed to monitor the compounds of interest^31^. Compounds were separated in a ZB-5plus capillary column (30 m x 0.25 mm x 0.25 μm, Agilent Technologies, USA). Agilent MassHunter Qualitative, Quantitative, and Unknown Analysis were used for data analysis. The retention times for different sugars were obtained with pure monosaccharide standards that underwent the same derivatization procedure.

### 2.7. Comparative homology analysis of *R. toruloides* and mapping of putative EPS biosynthesis and transport pathways

In our experiments, we deployed the *R. toruloides* CCT0783 strain, which was previously sequenced and deposited but without a functional annotation of the genome^5^. Therefore, we performed a functional annotation of this genome sequence and compared it with *R. toruloides* IFO0880^33^, *R. toruloides* NP11^34^, and *Rhodotorula glutinis* ZHK^35^ to identify homologies in genomic sequences, transcriptome expressions, and proteome relevant to the pathways investigated in this study.

#### 2.7.1. Functional annotation of *R. toruloides* CCT0783 genome

We deployed Braker3^36–48^, a fully automated genome annotation pipeline, to structurally annotate the genome of *R. toruloides* CCT0783, together with proteins of fungi compiled from OrthoDB database, offering evolutionary and functional annotations of orthologous genes with a wider sampling of genomes^49^. Braker3 was used with default parameters except for the addition of “—fungus” specifier. Reads were aligned to the CCT0783 genome using Hisat2 with a “—dta” parameter.

We utilized AnnoPRO^50^ for functional annotation of GO terms. The thresholds for filtering the outputs were 0.25 for biological processes, 0.15 for molecular functions, and 0.29 for cellular components. The thresholds were chosen based on a test run using *Saccharomyces cerevisiae* S288C proteome and functional annotation from yeastgenome.org^51^ to maximize the F1 score metric - a measure of the predictive performance of a model calculated based on precision and recall. CLEAN was used for the functional annotation of Enzyme Commission (EC) numbers. All EC numbers with confidence above 70% were added to the annotation^52^. The annotated genome of *R. toruloides* strain CCT0783 was then used to identify sequence similarities to the closely related reference genomes, namely (i) *R. toruloides* strain IFO0880 reference genome v4.0 (Mycocosm database)^33^, and (ii) *R. toruloides* strain NP11, taxid 1130832 (NCBI protein database)^34^. Julia, a programming language, was used to combine the outputs of Braker3, AnnoPRO, CLEAN, and the homology to NP11 and IFO880 into a single annotation^53^. Gene products were named based on NP11 names, IFO880 names, or names of corresponding EC numbers.

#### 2.7.2. Sequence similarity analysis of the annotated genome

We further focused on sequence similarity analysis of genes relating to EPS biosynthesis and transport using NCBI *blastp* command line software to find matches in the CCT0783 proteome for EPS and transporter genes from reference strains NP11 and IFO880, which, in turn, were obtained from UniProt^54^.

#### 2.7.3. Mapping the EPS biosynthesis and transport pathways, and the regulators of pH maintenance

A previous experimental study using *R. glutinis* ZHK yeast reported the presence of an EPS biosynthesis pathway and potential transporters for EPS export^35^. However, similar information is lacking for other *R. toruloides* strains, including CCT0783. For this reason, we performed the pathway mapping using the *R. glutinis* ZHK genomic dataset, which was retrieved from NCBI (genome assembly ASM1550198v1)^55^ and queried in a blast search against the reference genomes of *R. toruloides* IFO0880 and NP11 strains^33,34^. The default parameters (E=1e5, Filter=True, BLOSUM62) were used for the similarity analysis.

To complement the mentioned analysis and gain a system-level view of pathways, we queried genome-scale models of *R. toruloides*. These genome-scale models are based on genomic information from *R. toruloides* reference genomes and are used to stoichiometrically reconstruct enzymatic and transport reactions through gene-protein relation (GPR) information, which allowed us to query the genes corresponding to biochemical reactions in reference genomes. Genome-scale models of *R. toruloides* have been reconstructed using models of *Saccharomyces cerevisiae*, *Chlamydomonas reinhardtii*, human, mouse, *Escherichia coli*, and *Pseudomonas putida*, covering most of the orthologous proteins in *R. toruloides*^56^. We specifically queried the implicated pathway information from *R. glutinis* ZHK^35^ in the genome-scale models of IFO0880 and NP11 strains to identify potential genes involved in the synthesis of nucleoside diphosphate sugars (NDP sugars): ‘glucose 6-phosphate’ (g6p), ‘glucose 1-phosphate’ (g1p), ‘mannose 6-phosphate’ (man6p), ‘mannose 1-phosphate’ (Man1P), ‘UDP-glucose’ (UDP-glu), ‘UDP-galactose’ (UDP-gal), and ‘GDP-mannose’ (GDP-man) reactions^56,57^. Additionally, we utilized available information on EPS biosynthesis in other organisms, wherein gene candidates of glycosyl and mannosyl transferases are implicated in EPS production. Therefore, we searched publicly available omics data sets for *R. toruloides* strains using the keywords ‘glycosyltransferase’ and ‘mannosyltransferase’^7,56^, which allowed us to obtain the corresponding genes from the reference genomes.

Subsequently, we considered potential EPS transport candidates based on previous studies in other yeasts^8^. Here, we mapped two types of transporters: (i) within cell organelles, such as nucleotide-sugar transporters (NSTs) and intracellular proton transporters, and (ii) transporters on the cell membrane leading to the export of biopolymers. The transporters were searched from publicly available omics data sets using the keywords ‘UDP-’, ‘sugar’, ‘triose-phosphate’, ‘transporter’, ‘SLC’, ‘antiporter’, ‘ATPase’, ‘synthase’^7,56^. All the above allowed us to map genes in reference genomes, which were then used for sequence similarity analysis in the CCT0783 strain.

Finally, since several omics studies have been published on *R. toruloides* reference strains, including growth conditions, we used this data to compare the genes with high sequence similarity to the CCT0783 strain and to ascertain the presence of protein expression levels from relevant available omics datasets. Moreover, to better understand the synergy between nitrogen assimilation and the acidification of the CCT0783 culture, we also verified the presence of nitrogen transporters and ion exchangers that may be involved in maintaining intracellular pH and biopolymer production in this yeast.

## 3. Results

### 3.1. Understanding *R. toruloides* cellular aggregates

Our previous study of *R. toruloides* CCT0783 suspension cultures showed cellular aggregation in glucose-containing chemically defined medium^8^. In the study, we found that supplementing KCl and NaCl (0.1 to 1.0 mol/L) inhibited cellular aggregate formation^15^. However, the nature of extracellular biomolecules involved in cellular aggregate formation remained unexplored in the previous study. Hence, we decided to investigate possible extracellular macromolecules produced by *R. toruloides*. Here, we qualitatively and quantitatively identified and characterized extracellular macromolecules relating to aggregate formation. Firstly, we examined whether carbohydrate EPS were present in aggregate-inducing conditions since extracellular matrices in biofilms and microbial aggregates reportedly contain carbohydrates, proteins, and other macromolecules, depending on the nutrient environment^58,59^. For this, we deployed a qualitative method using the alcian blue stain^29^. This cationic dye binds to anionic polysaccharides through ionic interactions between its isothiouronium (-SC(NH_2_)_2_^+^) groups and deprotonated carboxylate (-COO^-^) or sulfate (-OSO_3_^-^) groups of the polysaccharides at low pH^60^. The stained samples, obtained by ethanol precipitation, showed a blue and cloudy matrix, indicating it is a carbohydrate-rich polymer (Fig. 1A). In this staining process, the phthalocyanine ring with a copper ion gives the blue color to the reaction product (Fig. 1B)^60^.

**Figure 1.**
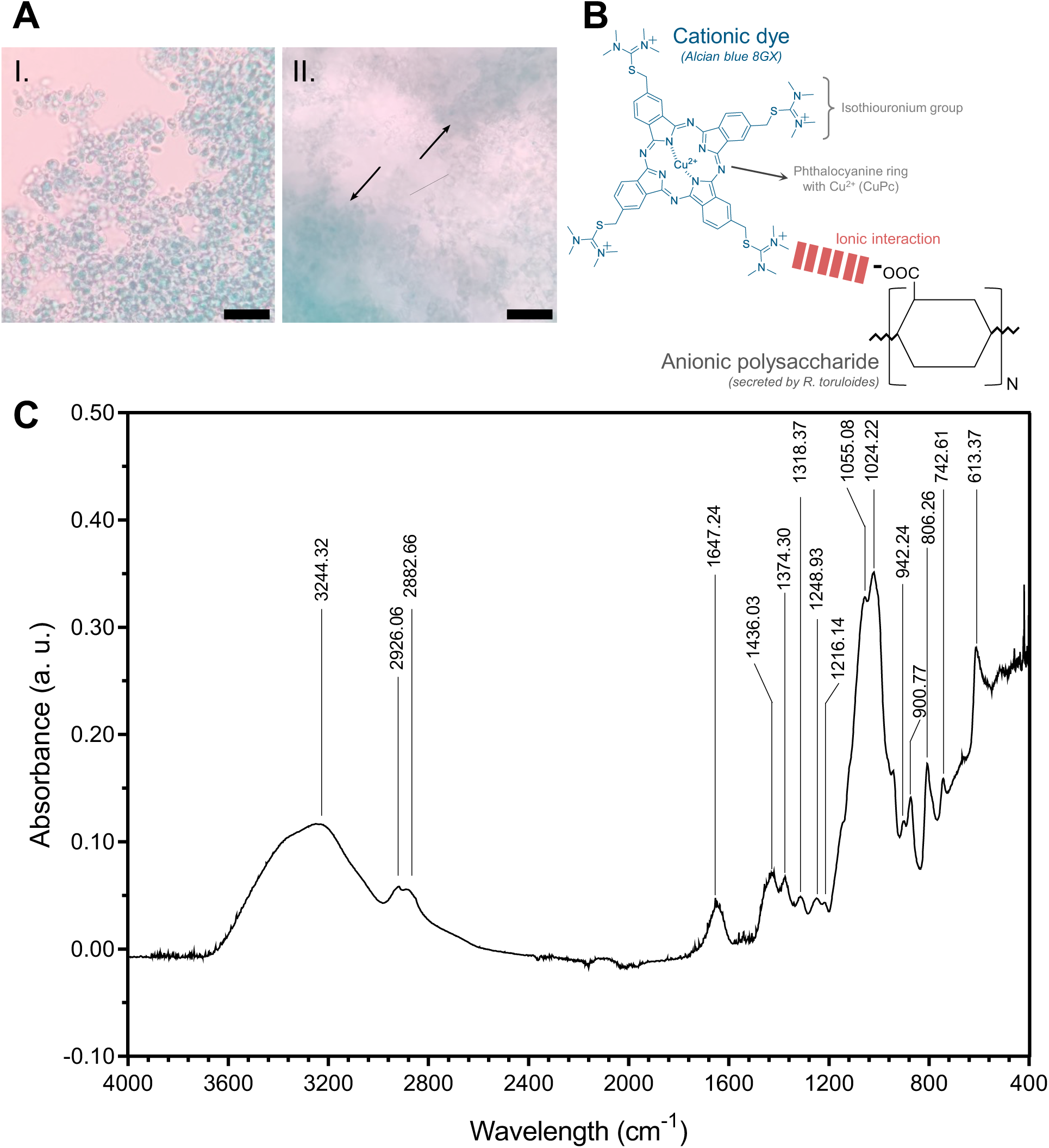
Characterization of *R. toruloides* cellular aggregates. **A)** EPS staining with alcian blue before (I) and after (II) precipitation with cold ethanol 96% v/v. Arrows point to precipitated EPS. Scale bar = 50 μm; **B)** Alcian blue stain interaction mechanism with anionic polysaccharides involves ionic interactions between charged groups; **C)** Representative FTIR spectrum of dried EPS.

Secondly, we ascertained the chemical fingerprints of the potential biopolymers in the crude extracts from *R. toruloides* cultures. To achieve this, we performed FTIR and GC-MS analysis on the dried EPS samples to determine characteristic peaks and chemical composition. The FTIR results showed characteristic peaks of carbohydrate polymers (Fig. 1C). The most significant peaks are presented in Table 1. The spectrum exhibits a broad peak for O-H stretching at 3244 cm^-1^. A double peak for aliphatic C-H stretching was observed at 2926 and 2882 cm^-1^. The peak at 1647 cm^-1^ corresponds to -O-H bending from water strongly bound to the EPS and/or C=O stretching. Combined with peaks at 1436 and 1374 cm^-1^, the EPS is likely acidic^35,36^. The ether peak for -CH_2_-O-CH_2_-was identified by two strong bands at 1055 and 1024 cm^-1^, usually between 1200 and 1000 cm^-1^. Peaks from 950 to 800 cm^-1^ indicate the presence of both α- and β-glycosidic bonds^61^. However, the peaks in the 1400 to 1200 cm^-1^ region could be attributed to impurities, including proteins and lipids in the crude EPS sample.

**Table 1.** - FTIR peak assignments for oven-dried EPS samples.

| Peaks (cm<sup>-1</sup>) | Assignments |
| --- | --- |
| 3244 | O-H stretching vibration |
| 2926, 2882 | Aliphatic C-H stretching vibration |
| 1647 | -O-H (H<sub>2</sub>O) bending vibration and/or C=O (carbonyl) asymmetrical stretching vibration |
| 1436, 1374 | C=O (carboxyl) symmetric stretching vibration |
| 1400 - 1200 | Impurities from proteins, lipids, and nucleic acids |
| 1200 - 1000 | C-O and CH<sub>2</sub>-O-CH<sub>2</sub> stretching vibration |
| 950 - 850 | α- and β-glycosidic bonds stretching vibration |

To determine the repeating units of the polysaccharide, the EPS was hydrolyzed using a TFA solution (2 mol/L), then reduced with a NaBH_4_ solution (20 g/L), and acetylated in a mixture of pyridine and acetic anhydride. GC-MS analysis of the treated EPS revealed that the monosaccharide composition of the carbohydrate fraction consisted of glucose (88.83 ± 4.87%), galactose (5.50 ± 1.50%), and mannose (4.80 ± 1.57%). Xylose and ribose were in a smaller quantity, pooling a total of 0.87 ± 0.04% of the carbohydrates (Fig. 2). The Bradford assay, using bovine serum albumin as the standard, indicated the presence of proteins. The assayed samples contained a protein fraction of 0.8% to 1.0% of its dry weight.

**Figure 2.**
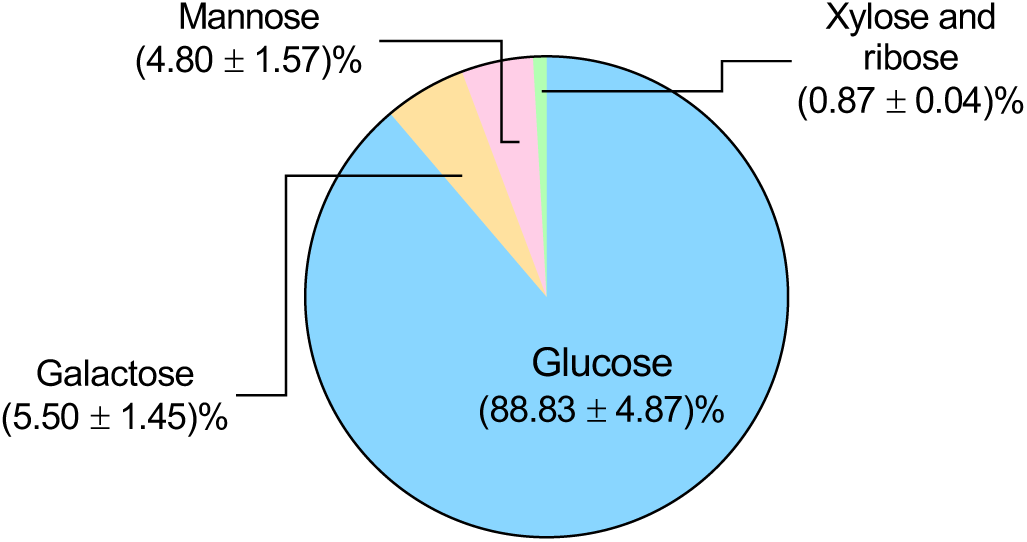
Monosaccharide composition of the carbohydrate fraction of EPS obtained through GC-MS analysis. The standard deviation corresponds to three independent experiments.

### 3.2. Evaluating EPS biosynthesis conditions

Having identified and characterized the EPS in conditions that enable cellular aggregate formation, we asked whether EPS is also produced during the suspension cultures’ cultivation, irrespective of aggregate formation (Fig. 3). We indeed found that *R. toruloides* CCT0783 produces EPS in suspension cultures, and not only when cells form aggregates (Fig. 3A). Next, we validated the role of potassium salts in the EPS production, which we previously reported in qualitative observation as prohibiting cellular aggregates formation in the suspension cultures^8^. Our results confirmed the previous study that supplementing cultures with additional potassium salt (KCl, 0.1 mol/L) caused them to remain free of cellular aggregates, without a significant reduction in biomass (Fig. 3A) However, we found that a higher ionic strength reduces EPS production by more than 50% in *R. toruloides* cultures, which likely contributes to the reduction in cellular aggregates.

**Figure 3.**
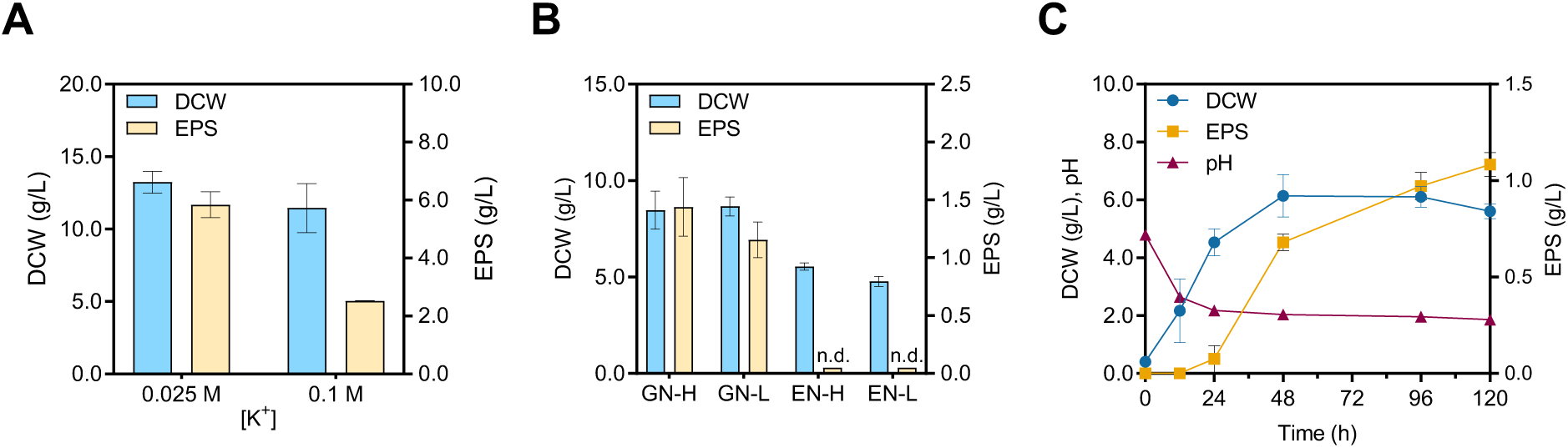
EPS production by *R. toruloides*. **A)** DCW and EPS (g/L) in chemically defined medium with glucose (120 g/L) and (NH_4_)_2_SO_4_ (5 g/L) supplemented with 0.1 M of KCl, compared to no KCl control, to increase [K^+^]. **B)** Variation in carbon source (glucose or ethanol) at 20 g/L and nitrogen source concentration (2.0 or 5.0 g/L) for dry cell weight (DCW) and exopolysaccharide (EPS) production quantification. GN-H: Glucose (2%), Ammonium sulfate (0.5%) - High; GN-L: Glucose (2%), Ammonium sulfate (0.2%) - Low; EN-H: Ethanol (2%), Ammonium sulfate (0.5%) - High; EN-L: Ethanol (2%), Ammonium sulfate (0.2%) - Low. n.d. means no EPS was detected during quantification. **C)** An example time series experiment showing DCW (g/L), pH, and EPS (g/L) graphs for cultivation up to 120 h in GN-L. The standard deviation corresponds to three independent experiments.

Subsequently, we investigated the potential metabolic route of biopolymer production in *R. toruloides*. For this, we cultured *R. toruloides* in glucose or ethanol-containing medium (Fig. 3B). We found that cultures grown in ethanol, a gluconeogenic carbon source, did not produce any detectable EPS but used this carbon for biomass. Moreover, the results showed that the carbon-to-nitrogen ratio in the culture media influenced EPS production. At 5 g/L of ammonium sulfate, an EPS titer of 1.44 ± 0.25 g/L was obtained (GN-H condition), while the titer of EPS was equal to 1.16 ± 0.15 g/L for 2 g/L of ammonium sulfate (GN-L condition) (Fig. 3B). In these experiments, the glucose concentration was kept constant at 20 g/L and the culture media initial pH was not fixed to a specific value and ranged from 4.40 to 4.57. We observed that the aggregate formation occurs during the early log phase for the conditions containing glucose with different nitrogen concentrations and dissipates after 24 to 48 h. We did not observe any cellular aggregation for ethanol-grown cultures.

Furthermore, we analyzed the temporal profile of EPS production over a 120-hour cultivation period for one of the EPS production conditions (namely, GN-L) (Fig. 3C). We observed that the EPS production occurs mostly during the transition from the growth phase to the stationary phase, reaching 1.08 ± 0.06 g/L after 120 h of cultivation. The biomass formation was associated with the acidification of the medium, from 4.79 to 2.18, in the first 24 h, during which only a fraction of EPS was produced. As the culture transitioned to the stationary phase, a further drop in the pH from 2.18 to 1.86 occurred, which corresponded with most of the EPS production (Fig. 3C). These results suggest that the glycolysis pathway is involved in the production of EPS. Therefore, in the subsequent experiments, we utilized glucose as the only carbon source and collected the EPS after 120 h for quantification.

### 3.3. Investigating the C:N ratio, nitrogen sources, and pH roles in EPS production

After observing the apparent role of glycolysis, we asked whether different carbon-to-nitrogen ratios in the culture media would affect EPS production. To achieve this, glucose concentration was increased from 20 to 120 g/L, and ammonium sulfate concentration was kept constant at 5 g/L in different triplicate batch culture experiments (Fig. 4A). We observed that the increased supply of glucose in distinct media cultures caused an increase in dry cell weight (DCW, 8.47 ± 0.99 to 13.2 ± 0.76 g/L) and in produced EPS (1.44 ± 0.25 to 5.84 ± 0.45 g/L).

**Figure 4.**
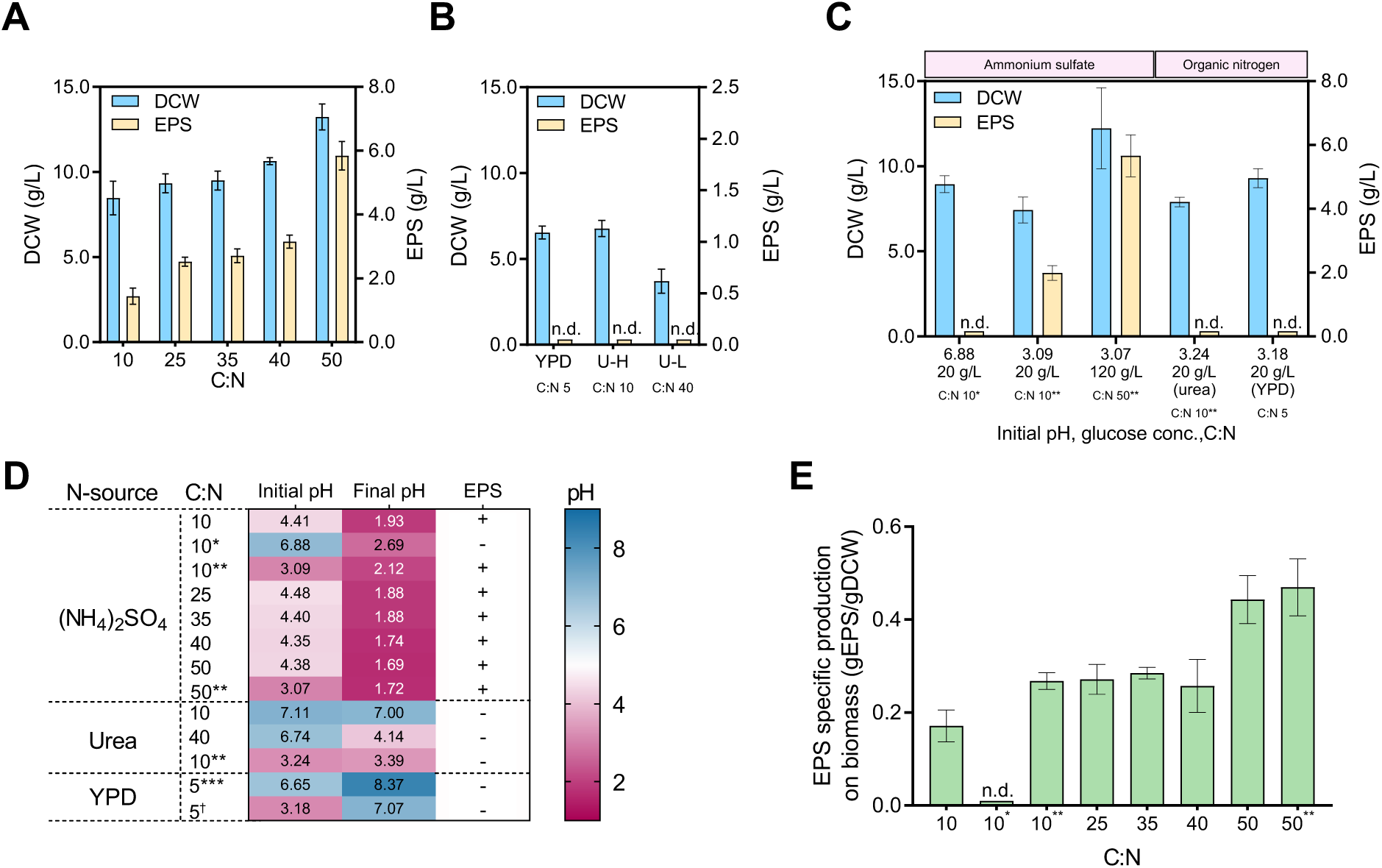
Evaluation of *R. toruloides* cultures in various culture media and cultivation conditions. **A)** Dry cell weight (DCW) and EPS (g/L) production at various C:N ratios, achieved by changing glucose concentrations (20, 60, 80, 100, and 120 g/L), while keeping (NH_4_)_2_SO_4_ constant (5 g/L) in the medium. **B)** DCW and EPS (g/L) in nutrient-rich medium (YPD) and chemically defined media with urea at 1.14 (U-L) or 2.28 (U-H) g/L as the nitrogen source. Glucose concentration was kept constant (20 g/L). **C)** DCW and EPS (g/L) in chemically defined media with glucose (20 g/L or 120 g/L) and (NH_4_)_2_SO_4_ (5 g/L) with phosphate buffer to pH 6.88, initial pH adjusted with HCl (1 mol/L) to 3.09 or 3.07. DCW and EPS (g/L) in chemically defined media with urea (2.28 g/L) and pH adjusted to 3.24. DCW and EPS (g/L) in YPD (20 g/L glucose) with pH adjusted to 3.18. **D)** Graphics indicating initial and final pH values, alongside the presence or absence of EPS in samples, are separated by the nitrogen source and C:N ratio. (*) Phosphate buffered condition with initial pH equal to 6.88. (**) pH adjusted with HCl (1 mol/L) to give an initial pH of around 3. (***) Estimated C:N ratio for YPD media is 5. (†) Estimated C:N ratio for YPD in pH adjusted with HCl (1 mol/L). (+) and (-) indicate EPS was detected or not detected, respectively. **E)** EPS specific production on biomass (gEPS/gDCW) after 120 h of cultivation in chemically defined media for different C:N ratios by changing the glucose concentration (g/L) of the media, where C:N 10 (20 g/L, pH not adjusted), 10* (20 g/L, pH adjusted to 6.88), 10** (20 g/L, pH adjusted to 3.09), 25 (60 g/L), 35 (80 g/L), 40 (100 g/L), 50 (120 g/L, pH not adjusted), and 50** (120 g/L, pH adjusted to 3.07). Only initial glucose concentrations were quantified. (NH_4_)_2_SO_4_ was kept constant (5 g/L). n.d. means no EPS was detected for quantification. The standard deviation corresponds to three independent experiments.

Further, since ammonium salt concentrations in our preliminary experiments appeared to influence EPS production (Fig. 3B), we examined whether organic nitrogen sources, such as urea (C:N ratios 10 and 40) in chemically defined media or a nutrient-rich YPD medium (approximate C:N ratio around 5), where peptone provides the nitrogen supply, would affect the EPS production. In those conditions, the yeast grew to reach DCWs of 6.53 ± 0.37 g/L in the YPD medium, and 3.70 ± 0.70 and 6.77 ± 0.47 g/L for U-L and U-H, respectively. A lower concentration of urea (U-L) caused a decrease in biomass compared to all other conditions with 20 g/L of glucose in the cultivation media. Interestingly, no EPS was detected (n.d.) in any of the organic nitrogen conditions investigated (Fig. 4B).

Our results suggest that nitrogen metabolism may play a role in influencing EPS biosynthesis (Fig. 4A, 4B). Previous studies have shown that inorganic and organic nitrogen sources uptake in yeasts involves distinct pH maintenance mechanisms^62,63^. Considering this, we further investigated EPS production at different pH conditions using both buffered and non-buffered media (Fig. 4C). For the inorganic nitrogen conditions, we compared a buffered medium with an initial pH of 6.88 to a non-buffered medium acidified to 3.09 with 1 mol/L HCl, while maintaining the glucose concentration at 20 g/L in both cases. The biomass and EPS formation differed between the buffered (DCW: 8.95 ± 0.49 g/L, EPS: not detected) and non-buffered cultivations (DCW: 7.43 ± 0.76 g/L, EPS: 1.99 ± 0.23 g/L), indicating that a lower starting pH in the latter reduced the biomass (12%, p-value < 0.1) and increased the EPS (38%, p-value < 0.05) produced compared with the non-adjusted pH condition (Fig. 4A, 4C). Moreover, for the condition with a glucose concentration of 120 g/L and an initial pH of 3.07, we observed DCW and EPS values of 12.23 ± 2.37 and 5.66 ± 0.65 g/L, respectively, which were not significantly different (p-values > 0.5) from the results obtained under non-adjusted conditions (Fig. 4A, 4C), suggesting a trivial role of the starting pH in increasing biomass or EPS production in this condition. The results on inorganic nitrogen potentially indicate a complex interplay between the C:N ratio and the initial pH of the media in biomass formation and EPS production. For the organic nitrogen sources, we tested pH-adjusted YPD (initial pH 3.18) and urea (initial pH 3.24) media containing 20 g/L of glucose to determine if a starting low pH would trigger EPS production; however, EPS was not detected in any of these conditions (Fig. 4C). Together, our results tentatively suggest that inorganic nitrogen metabolism, which is directly connected to extracellular pH drop^63^, is relevant to EPS production in *R. toruloides*.

To investigate a putative association between EPS synthesis and pH, we measured the initial and final pH values across all tested cultivation conditions (Fig. 4D). The media supplemented with ammonium sulfate, non-buffered conditions, and non-adjusted pH conditions had initial pH ranging from 4.35 to 4.48, which after 120 h of cultivation decreased to values at or below 2 in all tested conditions, and EPS was detected. Our findings indicate an apparent pH threshold for EPS synthesis under inorganic nitrogen-containing cultures. For the media with organic nitrogen, whether pH was adjusted or not, EPS was not produced. In urea-containing media, the final pH seemed to be dependent on the available urea in the media. The pH of the YPD media increased after 120 h, from 6.65 to 8.37, and from 3.18 to 7.17 in the pH-adjusted condition. The pH increase in YPD media is likely due to the consumption of amino acids, present in the rich media, and is consistent with previous studies^64^.

We summarized the specific production of EPS on biomass by calculating the gEPS/gDCW ratio for the cultures supplemented with ammonium sulfate (Fig. 4E). Among the conditions with a C:N ratio of 10, the culture with an initial pH of 3.09 produced 0.27 ± 0.02 gEPS/gDCW, whereas the condition starting at pH 6.88 did not produce detectable EPS. At higher C:N ratios of 25, 35, and 40, the specific EPS production values were 0.27 ± 0.03, 0.28 ± 0.01, and 0.26 ± 0.06 gEPS/gDCW, respectively. Finally, a maximum of 0.44 ± 0.05 gEPS/gDCW was observed at a C:N ratio of 50. When the same medium was acidified to an initial pH of 3, the specific EPS production increased slightly to 0.48 ± 0.06 gEPS/gDCW. Under this latter condition, a combination of a high glucose availability and an acidic environment suggests a limit to the EPS production of *R. toruloides* (Fig. 4E).

### 3.4. A comparative analysis of *R. toruloides* strains, focusing on a putative EPS biosynthesis pathway

To understand the potential metabolic pathways involved in the biosynthesis and extracellular biopolymer production in *R. toruloides,* we utilized available genomic, transcriptomic, and proteomic datasets. These datasets allowed us to perform a comparative analysis of *R. toruloides* strains, whose identifiable attributes were mapped for the CCT0783 strain, making its genome more accessible for further investigation. We retrieved the genome sequence of *R. toruloides* CCT0783 from NCBI^5^ and performed a functional annotation of this genome (Supplemental 1 or https://www.ncbi.nlm.nih.gov/bioproject/PRJNA1231390).

We used the annotated genome and data from previous studies to map and identify corresponding EPS biosynthesis and transport pathways. For this, we used available *Rhodotorula glutinis* ZHK experimental data that suggested an EPS biosynthesis pathway in this yeast^35^. Moreover, we categorized the existing *Rhodotorula* omics studies into two categories: potentially EPS-inducing conditions, where pH remained uncontrolled, and EPS-non-inducing conditions, where pH was controlled during cultivation (Table S1). In the absence of direct omics studies for the CCT0783 strain, this categorization enabled us to use homology and similarity searches to infer relative proteome changes in the putative EPS biosynthesis and transport pathways in EPS-producing and non-producing conditions (Table S1, Fig. 5).

**Figure 5.**
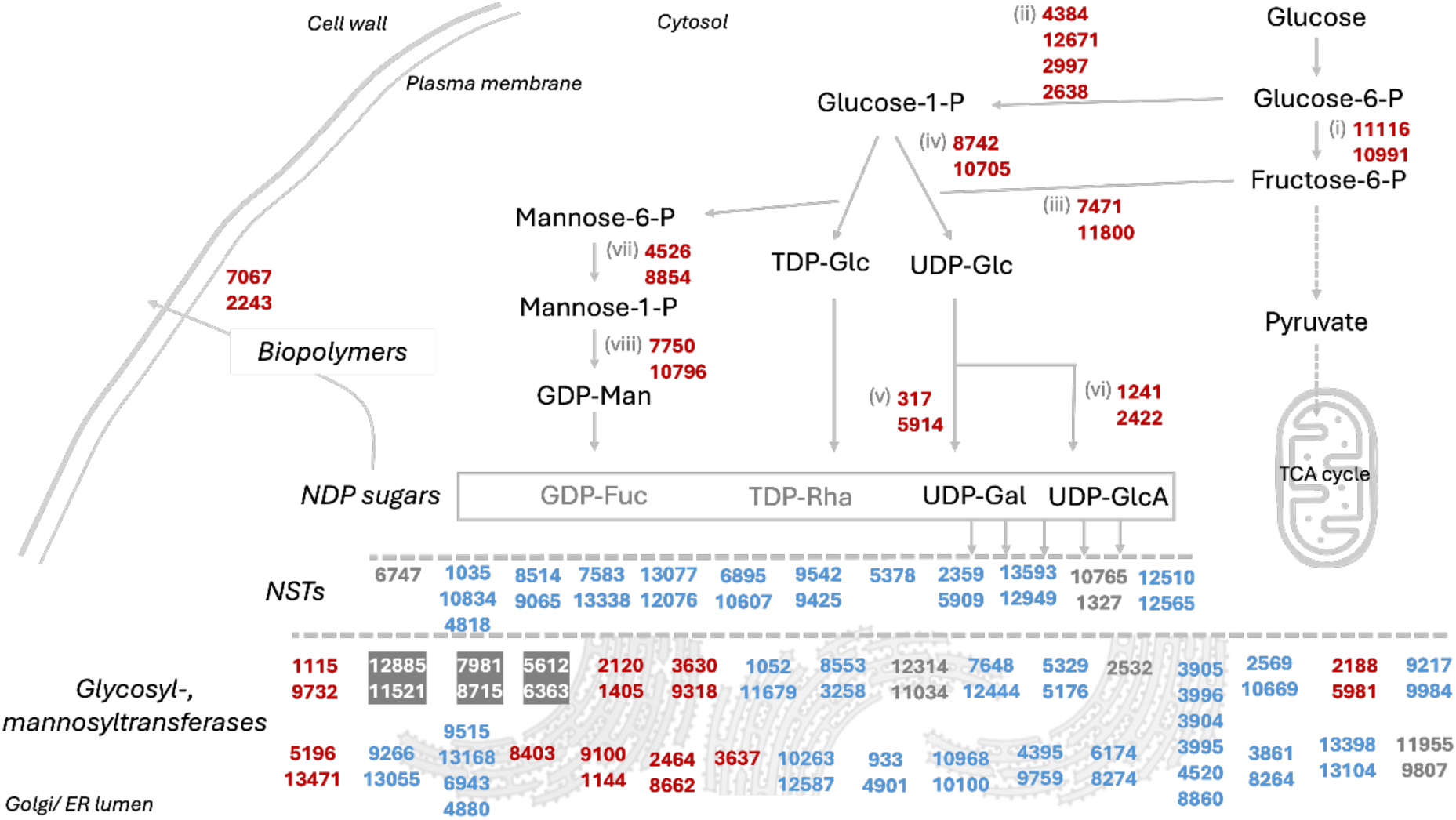
Proposed EPS biosynthesis and transport pathway in *R. toruloides* CCT0783, based on its genome annotation and representative gene expression data from previous omics studies. Names of genes, including reference studies, are listed in Supplementary Tables S2-S5. The mapped genes’ encoded proteins are visualized based on previous proteomics studies, highlighting differences in relative protein detection under EPS-inducing conditions: higher detection (red), lower detection (blue), and undetected proteins (gray). Proteins detected under control conditions are represented with a gray background. Abbreviations: (i) glucose-6-phosphate isomerase, (ii) phosphoglucomutase, (iii) mannose-6-phosphate isomerase, (iv) UTP-glucose-1-phosphate uridylyltransferase, (v) UDP-glucose-4-epimerase, (vi) UDP-glucose-6-dehydrogenase, (vii) phosphomannomutase, (viii) mannose-1-phosphate guanylyltransferase.

In a comparative gene sequence homology analysis with the reference *R. toruloides* strains (NP11 and IFO0880), CCT0783 strain genes encoding for NDP sugars (activated sugar donors for glycosylation reactions), NSTs (transport NPD sugars into the lumen of the Golgi or ER), glycosyl- and mannosyl transferases (catalyze the transfer of sugar residues from NDP sugars to growing polysaccharide chains), which are likely involved in the biosynthesis of EPS, showed 83% to 100% nucleotide similarity (Table S2, Fig. 5). We used this pathways sequence similarity information to estimate relative changes in the protein expressions for the EPS-producing and non-producing conditions (Table S2, Fig. 5). In the proteomics data, enzymes synthesizing NDP sugars are highly abundant in *R. toruloides*; however, neither upregulation nor downregulation on the protein level was observed by Kim et al.^56^ during EPS-inducing conditions (Table S3). In a comparison of exponential and stationary growth phases, Zhao and Li reported several NSTs upregulated in EPS-induced conditions in *R. glutinis* ZHK^35^. Therefore, to identify these genes in other *R. toruloides* strains, the gene sequences from *R. glutinis* ZHK were first retrieved from NCBI using annotations from Li et al.^55^ and then blasted against the genome of the reference *R. toruloides* strains. These Blast results identified several candidate genes with an identity score above 77% (Table S4). However, most NSTs were not identified or had very low abundance in the existing proteomics studies that used EPS-inducing conditions (Tables S4 and S5). This search also identified the glycosyltransferase RHTO_00642 with an identity score of 77.10% (Table S4), which was found to be abundant but downregulated under EPS-inducing conditions (Table S5). We identified several SLC35 genes in *R. toruloides* genomes, which encode a protein family responsible for transporting NSTs from cytosol or nucleus to ER or Golgi lumen^65^ (Table S5). The proteins encoded by those genes were identified as low abundant in *R. toruloides* in EPS-inducing conditions by Kim and colleagues^56^ (Table S5). However, some of them were not identified in the genome of CCT0783. NSTs are closely related to triose phosphate translocators (TPT) genes^66^, and we identified a few TPTs in low concentrations in EPS-inducing conditions (Table S5). Many mannosyl transferases responsible for glycosyl phosphatidylinositol synthesis in yeast are also present in *R. toruloides*^67^; some are highly abundant at proteome level (Table S5). Interestingly, the most abundant mannosyltransferases are downregulated during EPS production conditions but upregulated during the categorized control conditions. Based on the functional annotation of CCT0783 and a comparative homology analysis of available data, we propose a general pathway for the EPS biosynthesis in *R. toruloides* (Fig. 5). Though beyond the scope of the present study, we envision that a future experimental validation and analysis of the proposed pathway using the direct omics datasets for the biopolymers’ producing and non-producing conditions would be appropriate.

### 3.5. Unraveling the apparent role of pH in biopolymer production

Our experimental results suggest that inorganic nitrogen metabolism is associated with the acidification of the culture environment and simultaneous EPS production (Fig. 4D). A comparison between non-buffered and buffered media containing inorganic nitrogen indicates that EPS is produced when the final pH is around or below 2, which is observed in the former but not in the latter condition (Fig. 4). Moreover, organic nitrogen-containing cultures do not sufficiently acidify the media, and no EPS is detected. This indicates that *R. toruloides* assimilates inorganic and organic nitrogen sources through distinct mechanisms, as reported for other yeasts, where the nitrogen metabolism is linked with intracellular pH regulation^62,63^. Therefore, to unravel the pH maintenance processes in *R. toruloides* CCT07883, we performed a bioinformatics analysis of organellar and cellular membrane proton/ion and nitrogen transporters that likely contribute to the pH regulation (Fig. 6). In our analysis, firstly, three cation/H^+^ antiporters were identified to regulate intracellular pH in yeast, namely Nha1, Kha1, and Nhx1^68^, where Kha1 is localized in the membrane of the Golgi apparatus^69^. Secondly, low pH appears to increase the gene expression of ATP synthesis in an acidic environment^70^. The main regulator of pH in *S. cerevisiae* and other fungi is the P-type H-ATPase Pma1, which is present in the plasma membrane and is responsible for pumping protons outside of cells^71,72^. Further, V-type H-ATPases Vph1 and Stv1 are localized at the vacuolar membrane and between the Golgi apparatus and endosomes, contributing to pH maintenance^73,74^. We identified these genes in the CCT0783 strain, exhibiting 91% to 100% nucleotide similarity to those in other *R. toruloides* strains (Table S6). The proton/cation anti-transporters, except for the organellar transporters in vacuoles and mitochondria, were detected in low abundance or remained unidentified in EPS-inducing and control conditions in previous studies (Table S7). This could be attributed to the expected low abundance of membrane-bound proteins due to methodology limitations^75^. Interestingly, the organellar transporters Mdm38 and Vnx1 were not present in the data of the reference NP11 strain in a previous study^8^. However, one of the V-type H-ATPases (2783/3181) was highly abundant and upregulated during EPS-inducing conditions in earlier research^7,34,56^ (Table S7), indicating their role in the metabolism of *R. toruloides*. In yeast, protein kinase Sch9 (10750) has been shown to regulate V-ATPase assembly and disassembly, thereby controlling pH homeostasis in response to glucose availability^76^. We identified Sch9 in the genome of CCT0783 and found its presence in EPS-inducing conditions on the proteome level (Table S7). We also detected the gene sequences of Ca^2+^-ATPases Pmr1 and Ace2 in the CCT0783 genome, which have been reported to contribute to low pH adaptation in yeast^70^. Additionally, we identified two homologs of Pmr1 (13560/13419 and 6549/2724) that were previously observed at the proteome level under EPS-inducing conditions (Table S7). The genome also contains gene sequences encoding transporters for inorganic (ammonium permease) and organic (urea permease) nitrogen sources, exhibiting 89% to 100% similarity to other *R. toruloides* strains. These transporters likely play distinct roles in pH regulation and the induction of extracellular biopolymer production.

**Figure 6.**
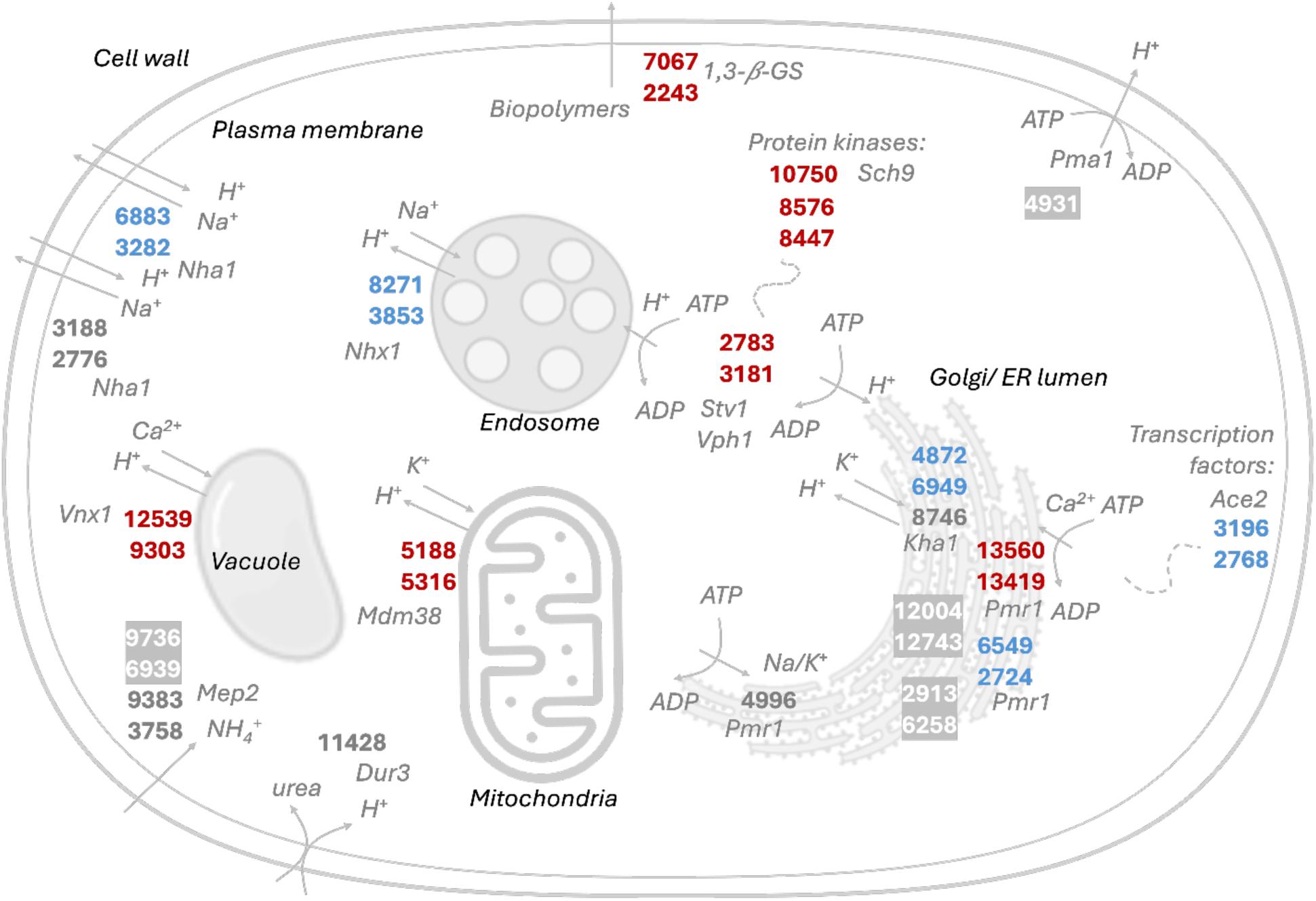
Putative proton, nitrogen, and biopolymer transport in *R. toruloides* CCT0783, based on its genome annotation and representative protein expression data from previous omics studies. Names of genes are listed in Supplementary Tables S6 and S7. The mapped genes’ encoded proteins are visualized based on previous proteomics studies, highlighting differences in relative protein detection under EPS-inducing conditions: higher detection (red), lower detection (blue), and undetected proteins (gray). Proteins detected under control conditions are represented with a gray background.

Finally, membrane-bound beta-glucan synthases have been associated with both the synthesis and transport of biopolymers, and we identified gene sequences related to these in the CCT0783 genome^77^ (Tables S2, S5-S7). Based on our analyses, we mapped putative intracellular proton transport, permeases, and biopolymer transport in *R. toruloides*, which may collectively contribute to intracellular maintenance of pH, the acidification of the culture media, and EPS production (Fig. 6).

## 4. Discussion

Our study systematically uncovers potential factors associated with extracellular biopolymer production in *R. toruloides* CCT0783, providing foundational results for conducting further mechanistic studies and engineering efficient biopolymer-producing cell factories. The experimental findings present insights into a biopolymer production process, chemically characterize EPS, and map the putative EPS biosynthesis and transport pathways.

Our qualitative analysis demonstrates that the produced biopolymer is mainly composed of polysaccharides, which was validated through FTIR and GC-MS analysis (Figs. 1, 2). The FTIR spectrum validated typical carbohydrate peaks previously described for microbial EPS produced by other *R. toruloides* strains^31,78^. Moreover, the dried EPS contained approximately 1.0% of proteins, indicating it is a complex biopolymer. Our results are consistent with previous reports for *Rhodotorula acheniorum* MC, in which 1.2% of the EPS was protein^23^. Pavlova et al. indicate that the crude EPS from *Sporobolomyces salmonicolor* AL_1_ contained 5.3% of proteins and 0.8% upon further purification^27^. Besides proteins at 0.8%, the EPS from *Rhodotorula mucilaginosa* sp. GUMS16 contained nucleic acids (0.7%)^31^. Other non-carbohydrate compounds, including glycoproteins and phospholipids, can also be found in EPS due to the non-specificity of the extraction process with ethanol^28^.

GC-MS analysis demonstrated that the monosaccharide repeating units of the carbohydrate fraction are glucose, mannose, galactose, xylose, and ribose (Fig. 2). We performed a comparative literature analysis of polysaccharides produced by *Rhodotorula* yeasts, which shows a wide variation in EPS composition and production amounts (Table S8). Briefly, 54 g/L of EPS was produced from *Rhodotorula bacarum* Y68 and was characterized as a pullulan^79^. *Rhodotorula* sp. strain CAH2 produced 7.5 g/L of EPS with sucrose as a carbon source, and it was composed of glucose, mannose, and galactose^80^. These three sugars are among the most reported components of microbial EPS in the literature, often accompanied by smaller proportions of arabinose, fucose, and rhamnose^81–84^ (Table S8). Some interesting examples of extracellular bioploymers include the EPS from *Rhodotorula acheniorum* MC that consisted of mannose (92.8%) and glucose (7.2%)^28^; when cultivated in co-cultures with yogurt starters, *Rhodotorula rubra* GED10 produced EPS that consisted mainly of mannose (83%), with small amounts of glucose, galactose, arabinose, and xylose^85^. In *Rhodotorula,* mannose is the most frequently reported monosaccharide in EPS, often accounting for up to 50% of the total monosaccharide content. It has been suggested that some resistance to antimicrobial substances and osmotic stress is associated with the ecological and physiological roles of EPS in yeast^28,86^.

Nevertheless, variations in strains and cultivation conditions play a significant role in determining EPS composition and, consequently, its properties. In our study, experiments using glucose or ethanol as carbon sources establish the role of glycolytic metabolism in EPS biosynthesis by *R. toruloides.* The use of the intermediates (glucose-6-phosphate and fructose-6-phosphate) of the upper part of the glycolysis pathway for EPS production is consistent with reports from other organisms, like *Lactococcus lactis*^87^. The growth and EPS production profiles for *R. toruloides* in our study are consistent with those of *R. mucilaginosa* sp. GUMS16 and other related yeasts^31^. Similar to our results, generally, no EPS is observed at the beginning of cultivations, and the maximum EPS production is shown during the stationary phase (Fig. 3). The C:N ratio of our experimental results demonstrates an increase in EPS production with higher glucose concentrations (Fig. 4A). However, this effect did not materialize when organic nitrogen sources (urea or peptone) were used (Fig. 4B, 4C). Moreover, EPS production is observed when the final pH approaches 2 in cultivations with an inorganic nitrogen source (Fig. 4D). No such EPS production is observed under cultivation conditions where the culture environment is buffered to neutral pH levels, or the pH remains above 2.5 (Fig. 4C, 4D). This observation is consistent with previous studies, for instance, Pavlova et al. explained the importance of the pH value to produce EPS by *Sporobolomyces salmonicolor* AL_1_, where the acidification of the medium during cultivation is a typical feature and a regulatory factor for EPS biosynthesis by this yeast^27^. Similar results were observed for *Candida famata* (*Debaryomyces hansenii*) and *Candida guilliermondii* (*Meyerozyma guilliermondii*) strains cultivated in media containing ammonium salts, in which EPS production occurred after the media pH dropped to 2.18 and 2.78, respectively^88^. Cho, Chae, and Kim reported that the maximum EPS titer (4.0 g/L) by *Rhodotorula glutinis* KCTC 7989 was achieved in the medium containing 2.0 g/L of ammonium sulfate with an initial pH of 4.0^89^. A potential mechanism of such acidification was proposed by Peña, Pardo, and Ramírez, where the consumption of ammonium salts by *Saccharomyces cerevisiae* was reported to decrease the pH of the culture environment^63^.

We performed a bioinformatics analysis to gain further insights and appreciate potential mechanisms involved in biopolymer production. This allowed us to map a potential EPS biosynthesis and transport pathway, alongside nitrogen transporters and ion exchangers involved in the pH maintenance of *R. toruloides* (Fig. 5, 6). Demonstrating the connection between pH and EPS production, our analysis identified the P-type H-ATPase Pma1 in the genome of strain CCT0783, showing 99.33% similarity to the NP11 strain. Its presence suggests that *R. toruloides* actively exports protons to the extracellular environment to maintain internal pH homeostasis, which is also observed in other yeasts (Fig. 6, Table S7)^72^, implying that the nitrogen metabolism-linked proton exchange by *R. toruloides* might be a major contributor to culture medium acidification.

In summary, glycolysis supplies both energy and precursor metabolites, some of which may contribute to the synthesis of UDP and NDP-sugars, which are key building blocks for the biosynthesis of extracellular biopolymers (Fig. 5). In parallel, nitrogen metabolism provides amino acids, balances the electrochemical potential in the cytosol, and contributes to the maintenance of intracellular pH in this yeast (Fig. 6). When inorganic nitrogen sources, such as ammonium, are consumed, proton pumps in the yeast ensure that the intracellular pH is maintained, causing the acidification of the extracellular environment, and EPS is observed concurrently. In contrast, the assimilation of organic nitrogen sources, like urea, involves different pH maintenance mechanisms that do not cause sufficient acidification of the environment nor produce EPS.

In conclusion, using our experimental results (Fig. 4) and bioinformatics analysis (Figs. 5,6), we propose that EPS biosynthesis is likely activated by medium acidification associated with nitrogen consumption. Overall, we expect our study to support further investigations of *R. toruloides*, developing this oleaginous yeast as a biopolymer cell factory in the future.

## Supporting information

Supplemental 1: Functional annotation of R. toruloides CCT0783 genome

Supplemental 2: Comparative analysis of R. toruloides strains

## Supplemental Data

Supplemental 1: Functional annotation of *R. toruloides* CCT0783 genome (attached file: CCT0783_annotation.gff3 or https://www.ncbi.nlm.nih.gov/bioproject/PRJNA1231390)

Supplemental 2: Comparative analysis of *R. toruloides* strains (attached file: Supplementary_tables_EPS.xlsx)

| Table | Title |
| --- | --- |
| S1 | Overview of experimental conditions. |
| S2 | Genes associated with EPS biosynthesis in *R. toruloides*. Identity refers to similarity check using Blast tool. |
| S3 | Gene expression studies associated with EPS production to synthesize NDP sugars in *R. toruloides* based on proposed pathway of EPS production in *R. glutinis* ZHK (Zhao and Li 2022) using the keyword search in the genome-scale models of *R. toruloides* IFO0880 (Kim et al. 2021) and NP11 (Tiukova et al. 2019) |
| S4 | Gene expression studies associated with EPS production in *R. toruloides* mapped from *R. glutinis* ZHK (Zhao and Li 2022) by using differentially expressed genes during EPS production |
| S5 | Gene expression studies associated with EPS production in *R. toruloides* mapped based on keyword search from a putative pathway of EPS production from various literature sources. |
| S6 | Genes associated with proton transport and biopolymer transport. Identity refers to similarity check using Blast tool. |
| S7 | Gene expression studies associated with intracellular proton and biopolymer transporters mapped based on Illarionov et al. 2021 and other studies |
| S8 | Quantification and characterization of EPS produced by *Rhodotorula/Rhodospiridium* yeast in the current literature |

## Acknowledgements

This work was supported by the Estonian Research Council (grant PRG1101) and from the European Union’s Horizon Europe Teaming for Excellence program (grant 101060066), which is co-funded by the Estonian state budget. KO acknowledges the funding and mobility grants from the Austrian government’s public employment service (AMS) and the European Research Council’s Erasmus+ program. PJ acknowledges the Estonian Center of Analytical Chemistry (ECAC), funded by the Estonian Research Council (TT4). BM and VP acknowledge support by the Swedish Research Council for Environment, Agricultural Sciences and Spatial Planning (Formas) (Grant Number 2018-01877).

Author’s contribution using CRediT standard: Writing – Original Draft Preparation (HSDRH, OT, AR, AI); Writing – Review & Editing (HSDRH, OT, KO, AR, AI, PJ, VP, PJL, RK); Conceptualization (HSDRH, AR, AI, RK); Methodology (HSDRH, OT, KO, AR, AI, PJ); Investigation (HSDRH, OT, KO, AR, AI, PJ, PMO); Data Curation (HSDRH, OT, KO, AR, AI); Software (AR, AI); Formal Analysis (HSDRH, OT, KO, AR, AI); Visualization (HSDRH, KO, AR, AI); Validation (HSDRH, OT, KO, AR, AI, PJ); Supervision (PJL, RK); Resources (PMO, GCMH, BM, NB, VP, PJL); Project Administration (VP, PJL, RK); Funding Acquisition (KO, PJ, BM, VP, PJL, RK).

